# Small Molecule Stabilization of PINK-1/PINK1 Improves Neurodegenerative Disease

**DOI:** 10.1101/2021.06.07.447442

**Authors:** Elissa Tjahjono, Jingqi Pei, Alexey V. Revtovich, Terri-Jeanne E. Liu, Alisha Swadi, Natalia V. Kirienko

**Affiliations:** Department of BioSciences, 6100 Main St, MS140, Rice University, Houston TX 77005 USA

**Keywords:** mitophagy, PINK-1, high-throughput, high-content, compound screening, neurodegenerative diseases, paralysis

## Abstract

Macroautophagic recycling of dysfunctional mitochondria, known as mitophagy, is essential for mitochondrial homeostasis and cell viability. Accumulation of defective mitochondria and impaired mitophagy have been widely implicated in many neurodegenerative diseases, and loss-of-function mutations of two regulators of mitophagy, PINK1 and Parkin, are amongst the most common causes of recessive Parkinson’s disease. Activation of mitophagy via pharmacological treatments may be a feasible approach for combating neurodegeneration. In this effort, we screened ∼45,000 small molecules for the ability to activate mitophagy. A high-throughput, whole-organism, phenotypic screen was conducted by monitoring stabilization of PINK-1/PINK1, a key event in mitophagy activation, in a *Caenorhabditis elegans* strain carrying a *Ppink-1*::PINK-1::GFP reporter. We obtained eight hits that induced mitophagy, as evidenced by increased mitochondrial fragmentation and autophagosome formation. Several of the compounds also reduced ATP production, oxygen consumption, mitochondrial mass, and/or mitochondrial membrane potential. Importantly, we found that treatment with two compounds, which we named PS83 and PS106 (more commonly known as sertraline) reduced neurodegenerative disease phenotypes (including delayed paralysis in a *C. elegans* Alzheimer’s model) in a PINK-1/PINK1-dependent manner. This report presents a promising step toward the identification of compounds that will stimulate mitochondrial turnover.

## Introduction

Although they are often simplistically characterized as the “powerhouse of the cell”, mitochondria have a wide range of cellular functions beyond that role, including amino acid metabolism, regulation of iron and calcium homeostasis, production of reactive oxygen species (ROS), stress surveillance, and control of apoptosis and other programmed cell death pathways (1-4). Considering the number and variety of roles that they play, it is not surprising that mitochondrial maintenance is crucial for the health of cells and organisms.

Unfortunately, cells have not many methods to repair mitochondrial damage. When damage is limited, mitochondria fuse, allowing their content to be mixed and sorted, segregating the damaged material (by an unknown mechanism) into low-quality mitochondria targeted for destruction via macroautophagic degradation (hereafter referred to as mitophagy) (5). The best-known pathway for triggering mitophagy is the PINK1/Parkin pathway, both of whose namesake members have been linked to Parkinson’s disease (5-7). Activation of this pathway begins with the PTEN-induced kinase 1 (PINK1), a serine/threonine kinase that is constitutively expressed and targeted to mitochondria (5). Upon arrival, healthy mitochondria will import the kinase and it will be destroyed by matrix-resident proteases. If mitochondria are damaged, PINK1 accumulates on the outer membrane, where it cross-phosphorylates itself, activating its ability to phosphorylate other substrates, including the E3 ubiquitin ligase Parkin. Once phosphorylated, Parkin will begin to ubiquitinate its targets, including outer mitochondrial membrane proteins, which allows their recognition by the machinery that recruits the isolation membrane and begins the process of engulfing mitochondria into an autophagosome. Once the autophagosome has closed, it will fuse with lysosomes to develop autophagolysosomes, where the contents will start to be degraded.

Mitochondrial turnover is activated under certain physiological conditions, such as during the maturation of erythrocytes (8), but evidence suggests that mitophagy participates in a broad variety of physiological functions, including in response to hypoxia and pathogen exposure (9-13). Additionally, mitophagy is surprisingly important for the maintenance of normal function for neurological cells. For example, mutations in both PINK1 and Parkin are linked with early-onset Parkinsonism (6, 14). Other types of mitochondrial dysfunction have been implicated in other neurodegenerative diseases (NDD), including Huntington’s disease, Alzheimer’s, and multiple sclerosis (7, 15-18), indicating failed mitochondrial maintenance as a common feature of these diseases.

Importantly, there is credible evidence that stimulating mitophagy can help mitigate some aspects of these disorders. For example, overexpression of PINK1 can restore mitophagy in a *Drosophila* model of Huntington’s disease and reduce disease symptoms (16). However, the consensus opinion is that treating most of these disorders will be more effective using small molecule therapies, rather than genetic modifications (19). Promising evidence in this field area exists as well. In an elegant recent study, Fang and colleagues demonstrated that a number of molecules, including urolithin A, could stimulate mitophagy and reduce disease burden in *Caenorhabditis elegans*, human neuronal cells, and two murine Alzheimer’s disease models (20). Additional efforts in this area have also been reviewed recently (19).

To date, a diverse set of *C. elegans* NDD models have been developed (21-23), and they have several valuable traits for this purpose, including a well-characterized nervous system, easy genetic manipulations, and highly conserved neurological pathways (24, 25). *C. elegans* is amenable for low-cost and high-throughput compounds screening due to its small size, short generation and lifespan, and the ability to simultaneously counterscreen for toxic compounds (24). Finally, as *C. elegans* is transparent, intact worms can be used for high-content screens where more nuanced or complex phenotypes can be used.

In this study, we used a high-throughput, high-content phenotypic screen using *C. elegans* carrying GFP tagged full-length PINK-1/PINK1 (*Ppink-1*::PINK-1::GFP) to identify a panel of small molecules that stabilize the kinase and verified their potentials in the activation of mitophagy. Four of the eight tested PS molecules significantly reduced mitochondrial membrane potential, while two of the others reduced mitochondrial mass loss. Neuroprotective properties of the selected PS compounds were tested in a transgenic *C. elegans* model of Alzheimer’s disease, where we discovered that two of the compounds significantly delayed paralysis in a PINK-1/PINK1-dependent manner. One of the two compounds was also able to reduce aggregate formation in a *C. elegans* polyglutamine-protein aggregation model. The compounds generally showed relatively low toxicity to human astroglial and prostate epithelial cells, increasing their promise.

## Results

### High-throughput screening of compound libraries identifies eight compounds that stabilize PINK-1/PINK1

As noted previously, preventing the degradation of PINK1 leads to activation of mitophagy (26-28) (**Fig. 1a**). To identify small molecules that promote PINK1 stability, we leveraged a *C. elegans* strain carrying a GFP-tagged, full-length PINK-1/PINK1 driven by its native promoter (9, 29). This reporter provides a simple method to track PINK-1/PINK1 stabilization by visualizing GFP. To develop a high-throughput, high-content phenotypic screen in *C. elegans*, we optimized parameters by identifying efficient conditions for the mitophagic activation by sodium selenite (Na_2_SeO_3_), which triggers the production of mitochondrial superoxide (30, 31) (**Fig. 1b**).

**Figure 1.**
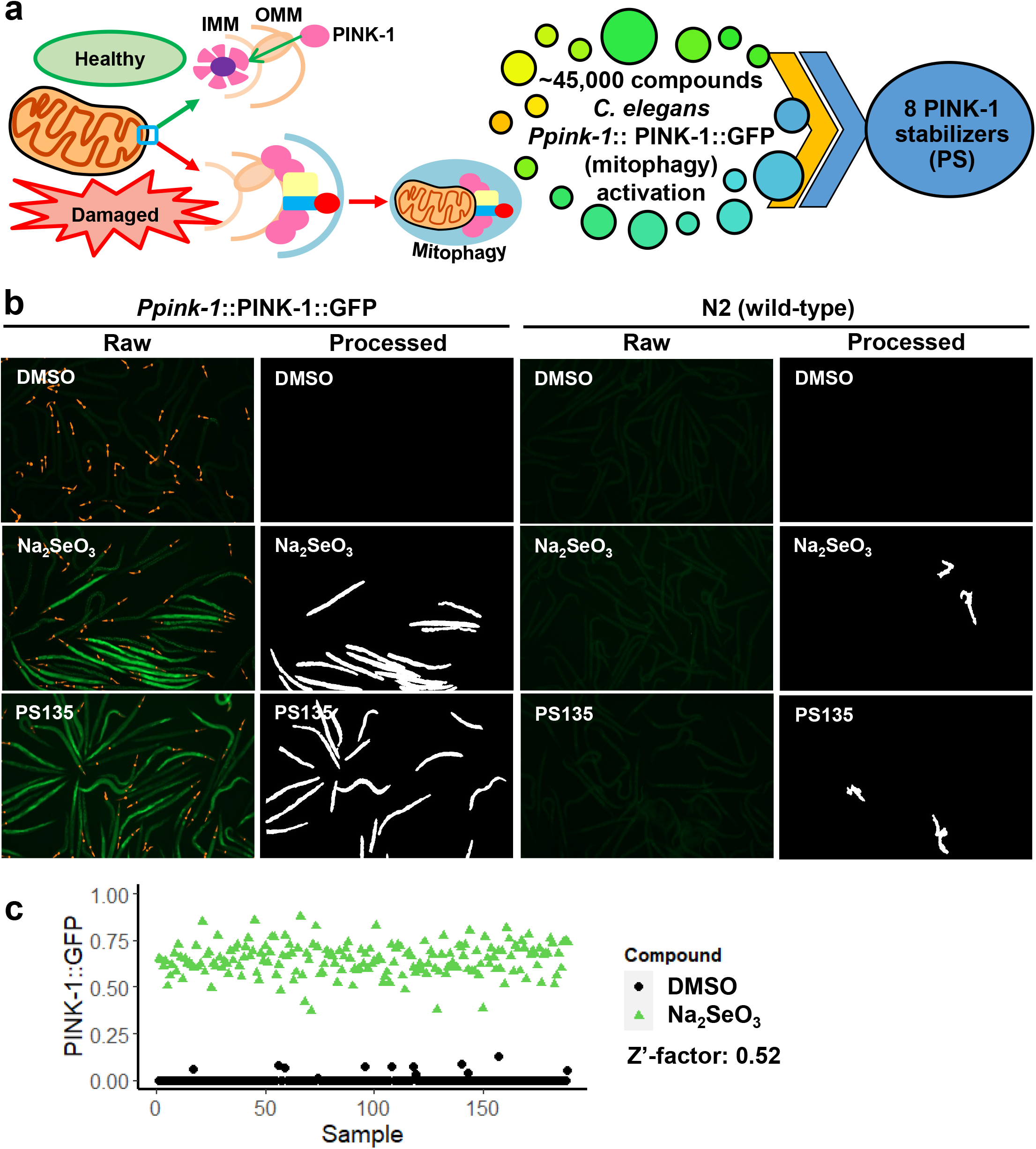
High-throughput, high-content PINK-1/PINK1 stabilization screen yielded eight hit compounds. **(a)** The accumulation of PINK-1/PINK1 in the outer mitochondrial membrane of damaged mitochondria initiates mitophagy. 45,000 compounds were screened in *C. elegans* resulting in eight PINK-1/PINK1 stabilizers. **(b)** Fluorescent (Raw) and Cell Profiler-processed (Processed) images of *C. elegans* carrying *Ppink-1*::PINK-1/PINK1::GFP or N2 wild-type upon 24 h of treatment with Na_2_SeO_3_, PS135, or DMSO control. Representative images are shown. **(c)** Quantification of GFP fluorescence of *C. elegans* carrying *Ppink-1*::PINK-1/PINK1::GFP after 72 h treatment with 7 mM Na_2_SeO_3_ or DMSO control. Three biological replicates for **(b, c)** were performed and analyzed.

Worms were exposed to the negative control or to sodium selenite for 24 h and then GFP was visualized using a Cytation5 multimode plate reader with automated process. GFP was quantified using an automated pipeline (to eliminate researcher bias) by using Cell Profiler software (32, 33) (Raw vs. Processed, respectively, **Fig. 1b**).

We determined the Z’-factor of the assay to be 0.52 (**Fig. 1c**). Z’-factors range from -∞ to 1, with the score indicating the degree of separation of the means and variability of the positive and negative controls, which indicates the ability of the assay to discriminate between signal and noise. A Z’-factor > 0.5 indicates a strong ability to detect even moderately weak hits.

Using this assay, we tested approximately 45,000 wells from various small molecule diversity and targeted libraries for stabilization of PINK-1/PINK1. Primary hit compounds were counterscreened in wild-type worms (which do not express GFP) to eliminate the possibility that the compounds themselves were fluorescent in worms. The eight compounds identified in this way were given the **PS** (for **<u>P</u>**INK-1/PINK1 **<u>S</u>**tabilizer) designations: PS30, PS34, PS83, PS103 (triclosan), PS106 (sertraline), PS127, PS135, and PS143 (**Fig S1, Table S1**). The final hit rate was 0.018%, which is somewhat lower than is commonly found from high-content screening in *C. elegans* (34, 35). Compound similarity analysis provided Tanimoto coefficients (36) that suggested substantial structural differences amongst the eight compounds (**Table S2**). This outcome was not surprising, as the hits primarily came from diversity libraries that were intended to explore broad amounts of chemical space.

### PS compounds trigger mitochondrial fragmentation and autophagosome formation

As the first step of characterizing the activity of the PS compounds on mitochondria, their impact on mitochondrial morphology was tested using a strain of *C. elegans* that expresses a mitochondrially-targeted GFP in body wall muscles (*Pmyo-3*::GFP^mt^). Under normal circumstances, these mitochondria take on a long, branched tubular network architecture that lies along muscle fibers. Worms treated with DMSO mostly showed this state (>85%, **Fig. 2a**)

**Figure 2.**
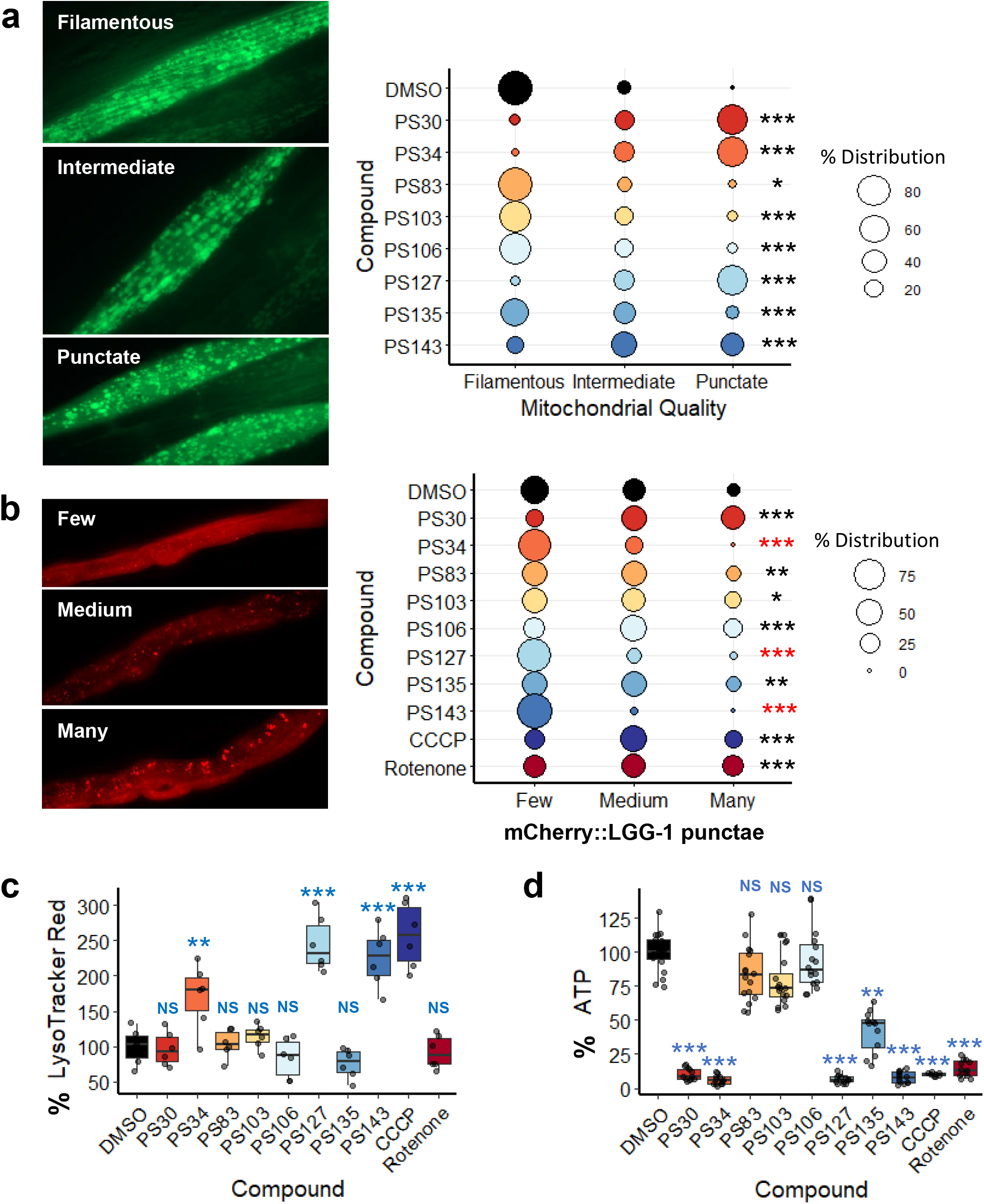
PS compounds induced mitochondrial fragmentation and mitophagy. **(a, b)** Fluorescent images and quantification of fluorescence of *C. elegans* carrying **(a)** *Pmyo-3*::GFP^mt^ or **(b)** mCherry::LGG-1/LC3 upon 15 h of treatment with PS compounds. **(c)** Quantification of LysoTracker Red fluorescence upon 15 h of treatment with DMSO, PS compounds, CCCP, or rotenone. **(d)** Quantification of luminescence (normalized to GFP) of *C. elegans* carrying *Psur-5*::luciferase::GFP upon 19 h of treatment with DMSO, PS compounds, CCCP, or rotenone. For **(a, b)**, percent distribution for each category was calculated and plotted, Chi-square statistic tests were performed, and representative images are shown. Three biological replicates with ∼30 worms/replicate were analyzed. For **(c, d)**, at least four biological replicates with ∼400 worms/replicate were analyzed. *p* values were determined from one-way ANOVA, followed by Dunnett’s test. All fold changes were normalized to DMSO control (at 100%). NS not significant, \**p* < 0.05, ** *p* < 0.01, *** *p* < 0.001.

Qualitative analysis of mitochondrial network structure showed that treatment with the PS compounds significantly increased the tendency towards intermediate (partially fragmented) or punctate (strongly fragmented) mitochondrial network. Four of the compounds, PS30, PS34, PS127, and PS143, induced substantial fragmentation, as shown by the dissolution of the tubular network into discrete punctae. PS83, PS103, PS106, and PS135 induced a milder disruption of the mitochondrial network, although each was statistically significant. Disruption of mitochondria fission-fusion homeostasis is consistent with existing evidence that mitochondrial fragmentation is necessary for mitophagy to occur (10, 37).

To confirm the formation of autophagosomes, we exposed a worm strain expressing mCherry::LGG-1 to the PS compounds. Under normal conditions, LGG-1, and its mammalian ortholog MAP1LC3, exhibit a diffuse cytoplasmic localization (38). During the formation of the isolation membrane (which will become the autophagosome), MAP1LC3 is crosslinked to the lipids that will comprise this membrane, converting the localization from a diffuse pattern into bright punctae (38, 39). Punctae were qualitatively assessed as few, medium, or many, with representative images shown (**Fig. 2b**). Compared to the DMSO control, several of the PS compounds increased punctae localization of mCherry::LGG-1/LC3, including two strong hits (PS30 and PS106) and three weak hits (PS83, PS103, and PS135), indicating increased autophagosomal formation. Intriguingly, three compounds (PS34, PS127, and PS143) reduced the production of autophagosomes, an unexpected outcome from stimulating the accumulation of PINK-1/PINK1. Two mitochondrial disruptors, rotenone (which blocks Complex I of the electron transport chain (ETC)) and carbonyl cyanide *m-*chlorophenyl hydrazine (CCCP, a proton uncoupler that dissipates the electrochemical gradient on the mitochondrial membrane), were used to validate the assay.

After formation of the autophagosome is completed, the next step in mitophagy is for the autophagosome to fuse with lysosomes to form autophagolysosomes (40). To observe this process, worms were treated with PS compounds, CCCP, rotenone, or vehicle control for 15 h. Afterward, worms were stained with LysoTracker Red DND-99, a dye that specifically labels acidic cellular compartments. This dye is routinely used to label lysosomes and autophagolysosomes (41, 42). Uptake was quantitatively measured by using flow vermimetry (43). Increased LysoTracker Red fluorescence was seen after treatment with PS34, PS127, PS143, and CCCP, but not with the rest of the PS compounds (**Fig. 2c**) (44). In contrast, treatment with PS30, PS83, PS103, PS106, or PS135 fragmented mitochondria but did not result in substantial formation of acidified organelles. This outcome was also seen for rotenone, which triggered the early stages of mitophagy (i.e., PINK-1/PINK1 stabilization, formation of the isolation membrane and autophagosome) but precluded acidification of the autophagosomes, preventing increased fluorescence from LysoTracker Red. This is consistent with reports elsewhere regarding the accumulation of autophagosomes and decreased autophagic completion after rotenone exposure (45, 46). This failure has been attributed to ATP depletion, which prevents the lysosomal vacuolar ATPase from consuming ATP to acidify the autophagolysosome (45).

Worms carrying a ubiquitously expressed firefly luciferase to provide real-time readout of ATP (47, 48) were treated with each of the eight PS compounds. Compounds that greatly induced mitochondrial fragmentation (PS30, PS34, PS127, and PS143, **Fig. 2a**) also caused significant drop in ATP production (**Fig. 2d**). Unexpectedly, there was no apparent correlation between ATP depletion and failure to acidify autophagolysosomes.

To investigate on whether ATP production failure was due to the inhibition of the ETC, we monitored the last stage of the chain by measuring oxygen consumption rate (OCR) (49). Only PS34, PS127, or rotenone significantly lowered oxygen consumption rate (with PS127 completely abolishing respiration) (**Fig. 3a**).

**Figure 3.**
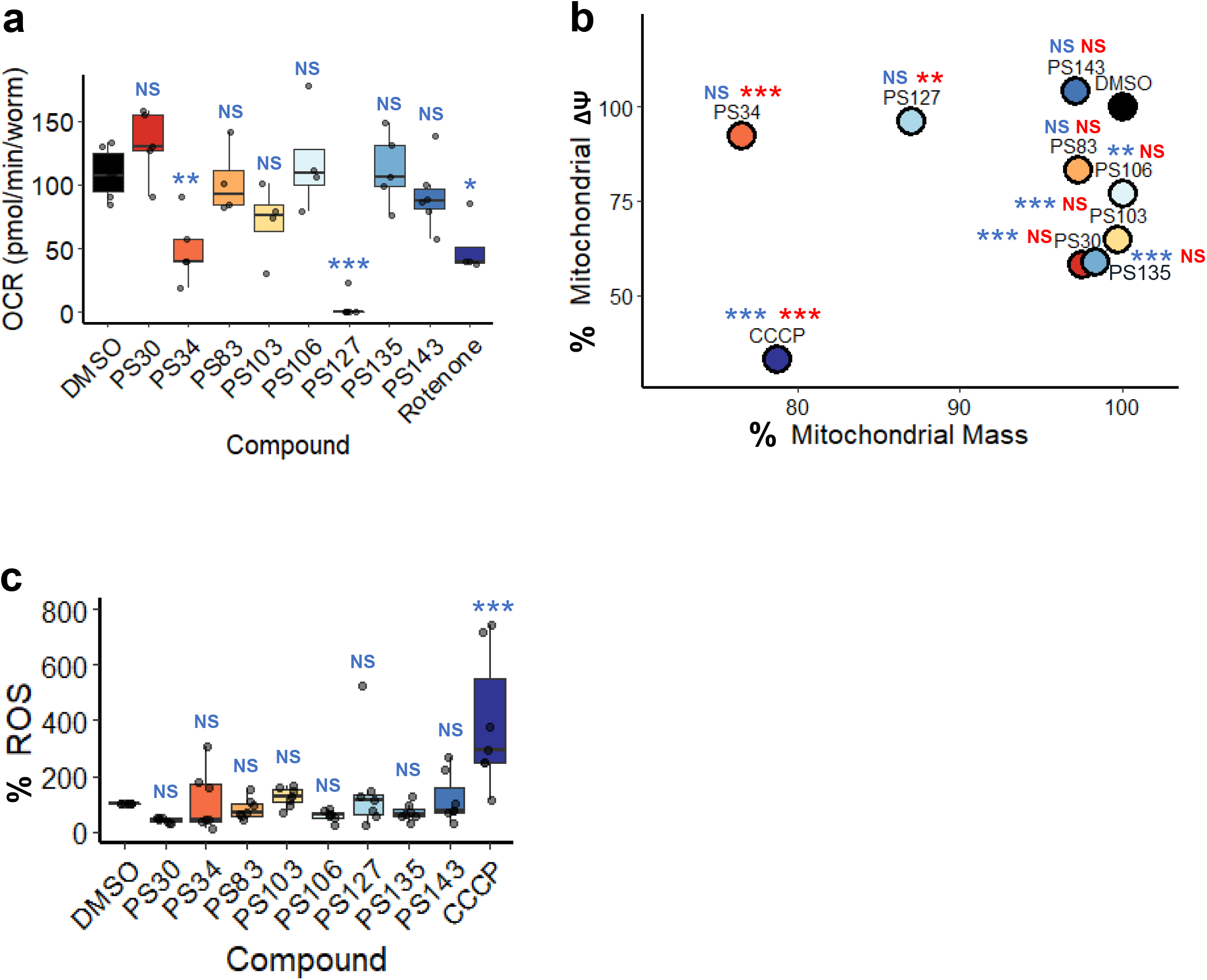
PS compounds differently affected multiple mitochondrial parameters. **(a)** Oxygen consumption rate (pmol/min/worm) measurement of wild-type worms upon 8 h of treatment with PS compounds or rotenone. **(b)** Point plot of nonyl-acridine orange fluorescence (mitochondrial mass, *x*-axis) and MitoTracker Red (mitochondrial membrane potential, *y*-axis) fluorescence of wild-type worms upon 15 h of treatment with PS compounds or CCCP. **(c)** Quantification of DHE fluorescence upon 15 h of treatment with PS compounds or CCCP. At least three biological replicates with **(a)** 6,000 worms/replicate or **(b, c)** ∼400 worms/replicate were analyzed. *p* values were determined from one-way ANOVA, followed by Dunnett’s test. For **(b, c)**, fold changes were normalized to DMSO control (at 100%). NS not significant, \**p* < 0.05, ** *p* < 0.01, *** *p* < 0.001. In **(b)**, blue color indicates significance of changes in mitochondrial membrane potential; red indicates significance of changes in mitochondrial mass.

### PS compounds show different effects on multiple mitochondrial parameters

To identify the mechanisms giving rise to mitophagy after compound treatment, several mitochondrial phenotypes were assayed. First, mitochondria were stained with MitoTracker Red, which accumulates in mitochondria proportionally to their membrane potential. Nonyl-acridine orange, a dye that binds to cardiolipin and is comparatively insensitive to mitochondrial membrane potential (50, 51), was used as a proxy to measure mitochondrial mass. CCCP, which dissipates mitochondrial membrane potential (52), was used as a control (**Fig. 3b**). CCCP reduced both mitochondrial mass and mitochondrial membrane potential, which is consistent with uncoupler treatment.

Significant reduction of mitochondrial membrane potential was observed in worms treated with PS30, PS103, PS106, and PS135 (**Fig. 3b**). This reduction did not appear to be accompanied by corresponding decrease in mitochondrial mass, may be indicative of weak uncoupling activity. These data suggest that for at least four compounds, the loss of mitochondrial membrane potential might be the trigger for mitophagy activation. In contrast, treatment with PS34 or PS127 reduced apparent mitochondrial mass, but not mitochondrial membrane potential, which may indicate that there are fewer mitochondria with increased membrane potential. We previously observed similar changes when worms’ diet was supplemented with vitamin B12, which improved mitochondrial health (53). Neither PS83 nor PS143 significantly affected mitochondrial membrane potential or mass (**Fig. 3b**). Combined, these results suggest that the loss of ATP content in PS30 and PS135 was not due to failure of the mitochondrial ETC, as in the case for PS34 and PS127. Instead, PS30, PS103, and PS135, and PS106 to a lesser extent, may have mild uncoupling activity.

Another common reason for the induction of mitophagy is the accumulation of ROS (54, 55). To test whether the PS compounds induced ROS production, worms were treated with compounds for 15 h, and then were stained with dihydroethidium, a non-fluorescent, redox-sensitive dye that is converted to fluorescent 2-hydroxyethidium by reaction with superoxide (56). Surprisingly, none of the compounds appeared to significantly increase ROS production (**Fig. 3c**). The positive control, CCCP, validated that the assay was being performed correctly (**Fig. 3c**).

### Four PS compounds trigger DAF-16/FOXO nuclear localization and SKN-1/Nrf2 pathway activation

Autophagic activation integrates a large number of signals, including stress and nutrient status. For this reason, we tested whether two master stress response pathways, the DAF-16/FOXO pathway and the SKN-1/Nrf pathway, are activated by exposure to the PINK-1/PINK1 stabilizing compounds. For DAF-16, a worm strain carrying a *Pdaf-16*::DAF-16a/b::GFP translational reporter (57) was treated with each of the compounds. Four compounds (PS34, PS83, PS127, and PS143) triggered DAF-16 translocation into the nucleus, demonstrating that it has been activated (**Fig. 4a**). The same four compounds also activated the conserved SKN-1/Nrf2 pathway, as shown by a transcriptional reporter (*Pgst-4*::GFP (58)) that is commonly used to confirm SKN-1 activation, albeit to a lower level than DAF-16 (**Fig. 4b**).

**Figure 4.**
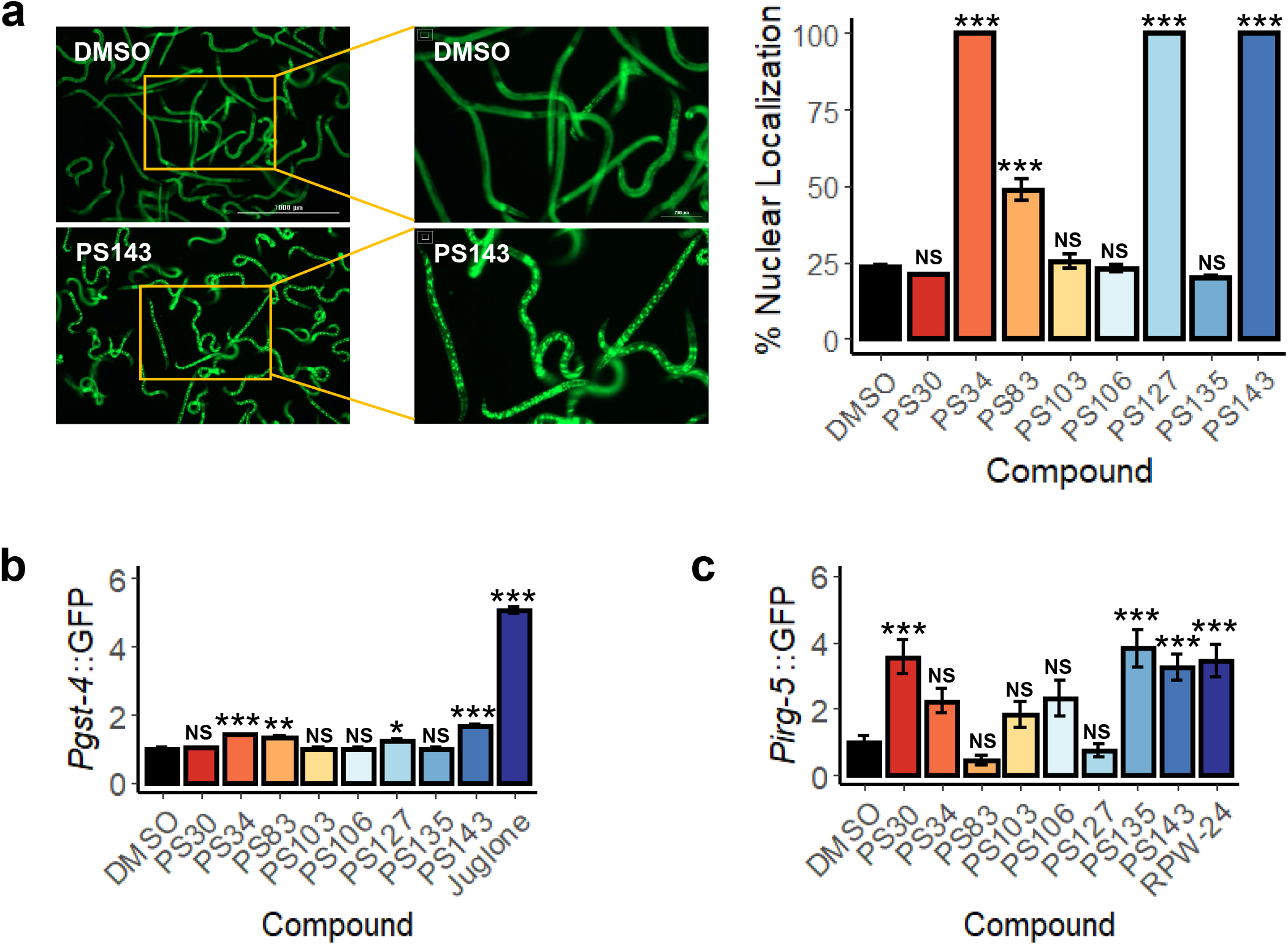
Four of the eight PS compounds activated DAF-16/FOXO and SKN-1/Nrf2 pathways. **(a)** Fluorescent images and quantification of nuclear localization (%) of *C. elegans* carrying *Pdaf-16*::DAF-16a/b::GFP upon 4.5 h of treatment with PS compounds. **(b, c)** Quantification of fluorescence of *C. elegans* carrying **(b)** *Pgst-4*::GFP or **(c)** *Pirg-5*::GFP upon 8 h of treatment with PS compounds. Representative images are shown in **(a)**. Three biological replicates with ∼400 worms/replicate were analyzed. Error bars represent SEM. *p* values were determined from one-way ANOVA, followed by Dunnett’s test. All fold changes in (**b, c**) were normalized to DMSO control. NS not significant, \**p* < 0.05, ** *p* < 0.01, *** *p* < 0.001.

Mitochondrial dysfunction that is sufficient to trigger mitophagy has also been shown to activate innate immune pathways (59-61). To test whether this occurred after treatment with these compounds, a worm strain carrying *Pirg-5*::GFP, a transcriptional reporter for the PMK-1 innate immune pathway (62), was exposed to the PS compounds or to RPW-24, a positive control. Only three of the compounds, PS30, PS135, and PS143, induced PMK-1 pathway activity (**Fig. 4c**).

### Two compounds, PS83 and PS106, show neuroprotective effects

Defects in mitochondrial clearance have long been linked to declined physiological function and NDD (as reviewed in (63)). On this basis, it is reasonable to hypothesize that the impacts of the compounds on mitochondria may stimulate protective effects in *C. elegans* models of NDD. A *C. elegans* model of Alzheimer’s disease was used to test this prediction. This strain, GMC101, produces full-length human β-amyloid in body wall muscles (64). Once the peptide has been expressed, shifting the strain to a higher temperature leads to β-amyloid aggregation and paralysis. Each compound was tested at 2-4 different concentrations (**Fig. S2**). Two compounds, PS83 and PS106, substantially reduced paralysis at 5 µM and 25 µM, respectively, and were comparable to the positive control metformin (**Fig. 5a-c and Fig. S2**). Another compound, PS103, provided a more modest, but still significant decrease in paralysis at 25 µM (**Fig. S2**). Alternatively, it is possible that PS83-and PS106-mediated rescue was an artifact and was independent of PINK-1/PINK1 stabilization. To test this, RNAi was used to knock down *pink-1* expression in the β-amyloid-expressing strain prior to compound exposure. Consistent with our interpretation, *pink-1(RNAi)* completely removed the ability of PS83 or PS106 to delay paralysis, and made worms treated with PS83 or PS106 indistinguishable from vehicle controls **(Fig. 5d, e)**.

**Figure 5.**
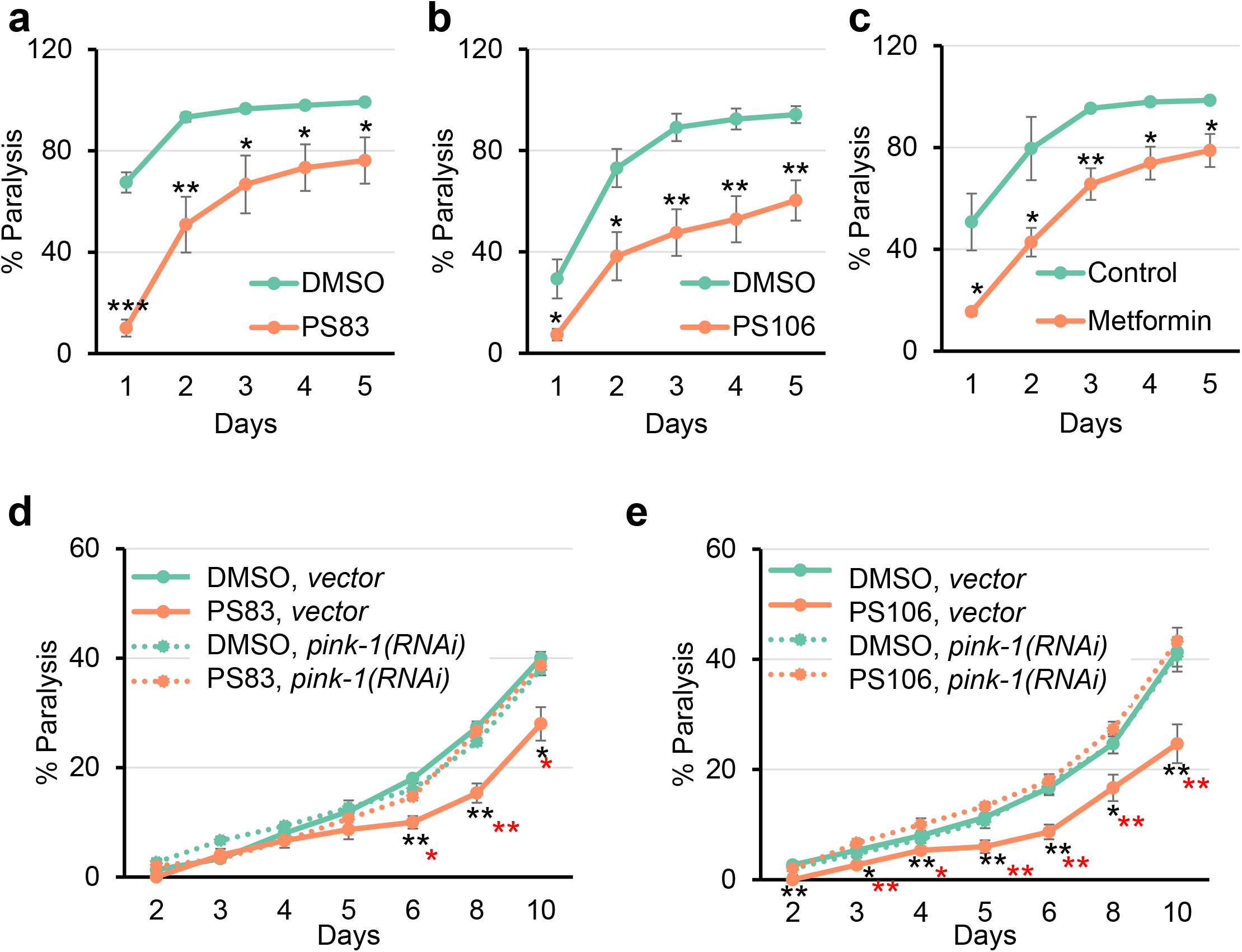
PS83 and PS106 reduced the rate of paralysis in *C. elegans* strain expressing human beta-amyloid. **(a-e)** Rate of paralysis curve of *C. elegans* Alzheimer’s model (GMC101) expressing full-length human beta-amyloid upon treatment with **(a, d)** 5 µM PS83, **(b, e)** 25 µM PS106, or **(c)** 100 mM metformin control. In **(d, e)**, worms were reared on *E. coli* expressing *cdc-25*.*1(RNAi)/vector(RNAi)* or *cdc-25*.*1(RNAi)/pink-1(RNAi)*. At least three biological replicates with ∼180 worms/replicate were analyzed. *P* values were determined from Student’s *t*-test. \**p* < 0.05, ** *p* < 0.01, *** *p* < 0.001. Black stars indicate significance compared to DMSO control. Red stars in (**d, e**) indicated statistical significance between PS compound on *vector* vs *pink-1(RNAi)*-reared animals.

To test whether the compounds had a broad effect on NDD models, we obtained a worm strain that expresses a YFP-tagged protein with an engineered polyglutamine (polyQ) repeat of 82 consecutive glutamine residues (Q82::YFP) (65). PolyQ repeats are causative for at least ten different neurodegenerative diseases, with the best-known being Huntington’s chorea (66). Expression of the chimeric Q82::YFP product under the control of a tissue-specific promoter (e.g., *unc-54* or *vha-6*) causes Q82::YFP aggregation in the target tissue (65).

Young adult worms expressing the Q82::YFP construct under the intestinal promoter *vha-6* were treated with PS83 (5 µM) or PS106 (25 µM) for 24 h, and then aggregates were manually counted under low magnification (**Fig. 6**). We found that PS83 showed no clear difference from vehicle alone (**Fig. 6a**), while PS106 treatment significantly reduced the number of aggregates (**Fig. 6b**). Previously, the Morimoto lab implicated activated DAF-16/FOXO in limiting Q82 aggregation and paralysis when aggregates form in body wall muscles (65). Since PS106 did not induce DAF-16 nuclear localization, the precise regulatory pathway induced by PS106 and its role in providing neuroprotection will need to be further elucidated.

**Figure 6.**
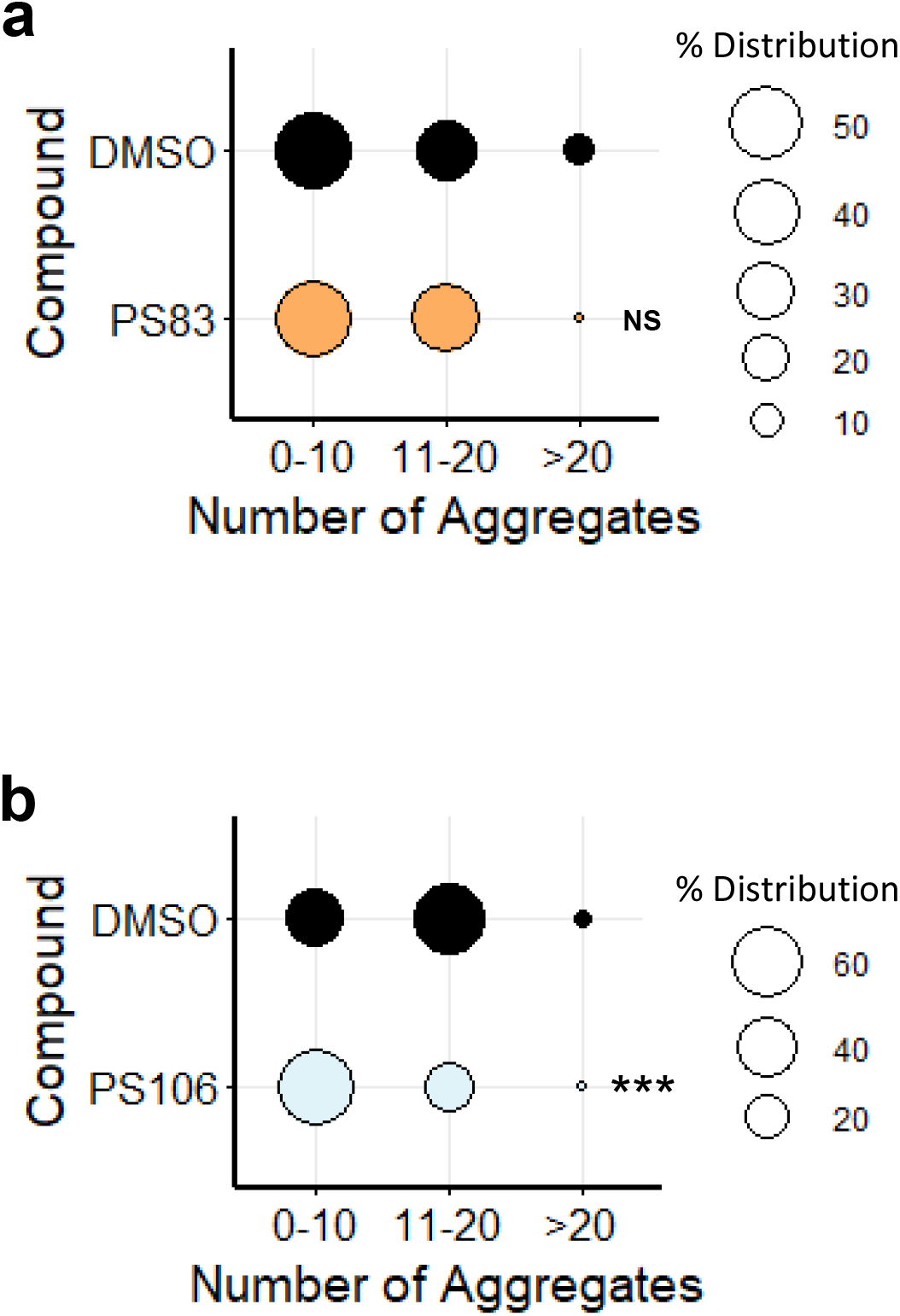
PS106 reduced aggregate formation in *C. elegans* strain expressing polyglutamine. **(a, b)** Percent distribution of the number of polyglutamine aggregates in *C. elegans* strain expressing polyglutamine (Q82) (GF66) upon treatment with **(a)** 5 µM PS83 or **(b)** 25 µM PS106. Three biological replicates with ∼30 worms/replicate were scored and analyzed. *p* values were determined from Chi-square test. NS not significant, *** *p* < 0.001.

### PS compound exposure shows limited toxicity

While activation of mitophagy can provide a means to overcome some aspects of NDD, overactivation of mitophagy may lead to excess mitochondrial loss, bioenergetic deficits, and cellular death (67). To test whether prolonged exposure to PS compounds causes death, survival of *C. elegans* and human cells was measured. For *C. elegans*, worms were exposed to the eight PS compounds at four different concentrations for 72 h and then were incubated with Sytox Orange, a cell-impermeant dye that stains DNA in dead worms. Treatment with most compounds showed greater than 75% of survival (as normalized to the DMSO control) at concentrations of 25 µM or less (**Fig. 7a**). In all cases, worms survived the lowest tested dosage; in all but one (i.e., PS83), the highest dose caused at least partial death (**Fig. 7a**). This suggests that some optimization would be necessary to see protective effects. Importantly, the two compounds that reduced paralysis rate in the *C. elegans* Alzheimer’s model (PS83 and PS106) did not impair survival at tested concentrations of up to 100 µM.

**Figure 7.**
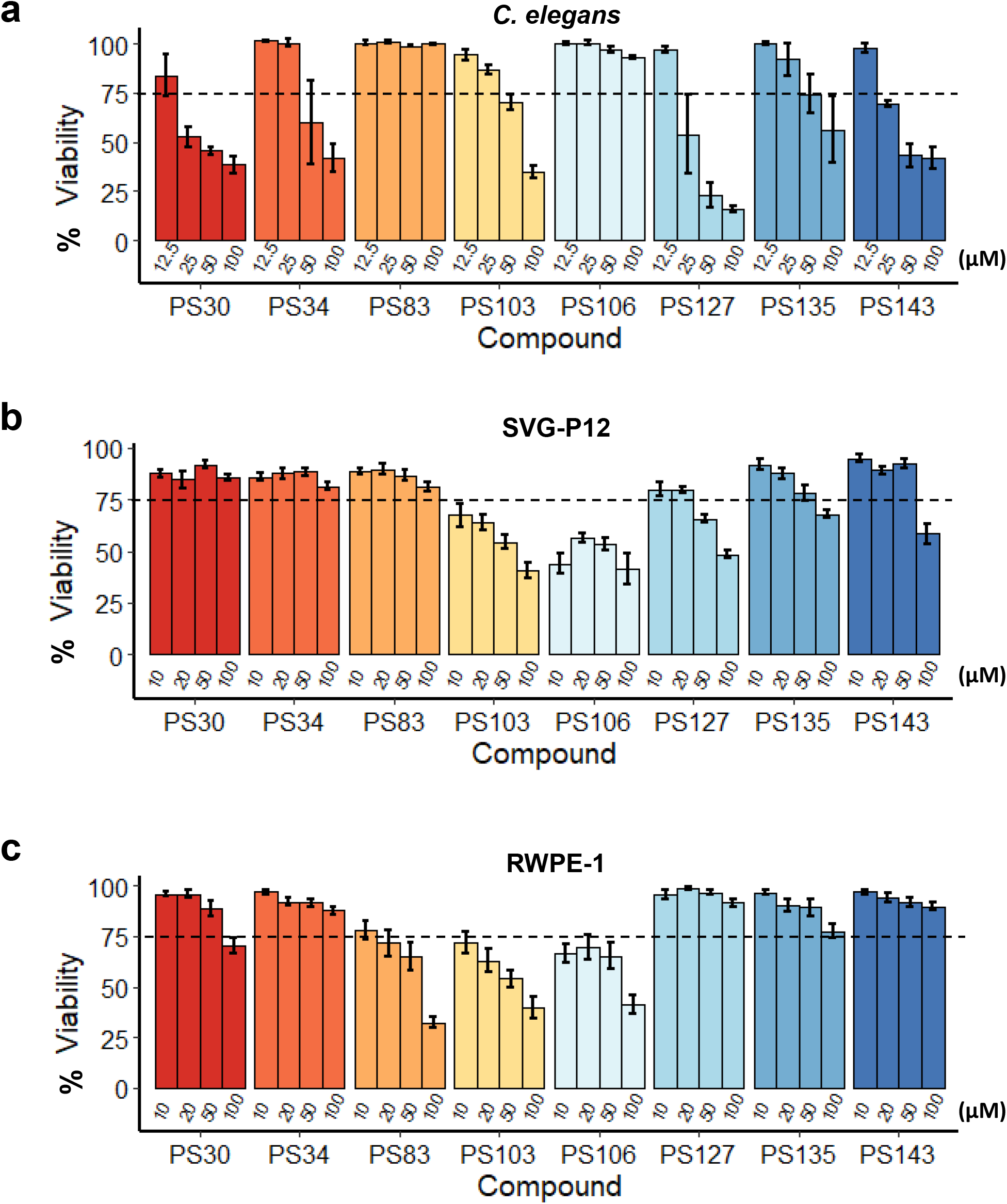
Most PS compounds have low toxicity in *C. elegans* and mammalian cells. Viability of **(a)** *C. elegans*, **(b)** human astroglial (SVG-P12) cell line, or **(c)** human prostate epithelial cell line (RWPE-1) upon 3 days of exposure to PS compounds with concentration gradient as indicated on the graphs. Dotted lines mark 75% of viability. At least three biological replicates were analyzed. Error bars represent SEM. All fold changes were normalized to DMSO control.

Cytotoxicity of chronic, 72 h exposure to PS compounds at similar concentrations was also measured in two human cell lines, SVG-P12 (an astroglial cell line) and RWPE-1 (prostate epithelial cells) (**Fig. 7b, c**). Differential Hoechst / propidium iodide staining was used to assess cell death. Exposure to either PS103 or PS106 at concentrations higher than 10 µM showed substantial toxicity in both cell lines. Interestingly, PS34, PS127, PS135, and PS143 were less toxic to human cells than to *C. elegans*.

## Discussion

Stimulation of mitophagy has proven to be a promising therapeutic target for neurodegenerative diseases (68) and may be beneficial for healthy aging. Using a high-throughput, high-content phenotypic screen, we obtained and characterized eight PINK-1/PINK1-stabilizing compounds. Interestingly, two of the compounds had previously been associated with alterations in mitochondrial function.

PS103, commonly known as triclosan or irgasan, has been linked with a variety of mitochondrial dysfunction, including uncoupling of the mitochondrial membrane potential by reversible protonation of the phenoxy group (69-71) and inhibition of Complex II of the ETC (72). Triclosan has also been associated with increased mitochondrial ROS, reduced mitochondrial mass, and disruptions in mitochondrial morphology (73). The ability of triclosan to cause several types of mitochondrial damage, apparently with different proximal factors (72), somewhat reduces its value as a therapeutic agent.

Despite its common appearance in a wide variety of consumer products, questions about the safety of triclosan remain, even if used externally. For example, in addition to its role in mitochondrial disruption, triclosan also has the potential to disrupt endocrine function, affect immunity, disrupt calcium and zinc homeostasis, and alter lipid metabolism (74). Triclosan also has a strong potential for bioaccumulation (75), which is undesirable in a maintenance medicine. Given these caveats, the potential for triclosan to be developed into a treatment for NDD seems small.

The potential for PS106, more commonly known as sertraline, is substantially greater. Sertraline is thought to bind to the serotonin transporter (SERT) in the presynaptic neuron, preventing reabsorption of serotonin and prolonging synaptic signaling. Sertraline is a well-known compound with carefully studied pharmacological effects and is one of the most commonly prescribed psychiatric medications in the US. Our data contribute to an ongoing discussion about the potential for sertraline as a treatment for one or more NDDs.

Probably the earliest hint that sertraline may have some unexpected effect on mitochondria came in a report attempting to identify ‘hidden’ drug targets, where the authors determined that sertraline had several characteristics similar to the well-known mitochondrial toxin rotenone (76). Not long after, Kumar and colleagues demonstrated that sertraline treatment could ameliorate damage caused by the mitotoxic agent 3-nitropropionic acid (77). Recently, it was demonstrated that sertraline prevents the function of the mitochondrial VDAC1, reducing cellular ATP, increasing the ADP/ATP ratio, and activating autophagy through mTOR (78).

Sertraline increases survival and neurogenesis at pharmacologically relevant concentrations in several murine models of Huntington’s (79, 80) and physiological outcomes (e.g., grip strength, coordination, locomotor activity, etc.) in rat models of Huntington’s and Parkinson’s diseases (81, 82). Given the frequent co-occurrence of depression with NDD, it is not surprising that sertraline is often prescribed to these patients suffering from these disorders. Promisingly, some Parkinson’s patients receiving sertraline have shown improvement in their symptoms (83, 84). It is clear, given these findings, that a more systematic study of the potential for sertraline for the treatment of NDD is warranted.

The remaining six compounds are considerably less well characterized. PS83, formally known as [4-[N-[(*E*)-2-cyano-3-oxo-3-thiophen-2-ylprop-1-enyl]-S-methylsulfonimidoyl]phenyl]4-chlorobenzoate, has several similarities in treatment outcome to sertraline and triclosan. For example, all three compounds caused relatively minor mitochondrial fragmentation, but little other effect. PS103 and PS106 caused greater depolarization, while PS83 apparently did not. Searching the literature for other reports of PS83 failed to provide additional clues. For the time being, we can only conclude that it stabilizes PINK-1::GFP and that this stabilization does not appear to be a consequence of depolarizing the mitochondrial membrane or disrupting the electron transport chain.

Two of the compounds, PS30 and PS135, appear likely to be mitochondrial uncouplers. They reduced mitochondrial membrane potential and decreased ATP content, but oxygen respiration continued unabated. Like PS83, relatively little is known about PS30 or PS135. However, an analog of PS30, known as SMTC1100, has been shown to be helpful in Duchenne muscular dystrophy (85). This fatal, progressive disorder is characterized by wasting muscle loss due to disruption of the dystrophin protein, which leads to mitochondrial dysfunction (86, 87). This suggests that a larger portion of the scaffold may have a positive effect on mitochondrial recycling in chronic degenerative disorders.

The final group of three compounds, PS34, PS127, and PS143 share a number of characteristics that indicate that they may be acting the same way. They show substantial mitochondrial fragmentation, autophagolysosomal acidification, they reduced ATP and oxygen consumption, and considerably reduced average mitochondrial mass. The compounds did not, however, reduce mitochondrial membrane potential, suggesting that they disrupt degradation of PINK-1/PINK1 in a different fashion. Interestingly, they were also amongst the strongest activators of GST-4/Nrf and DAF-16/FOXO, which may indicate that they are causing other damage to the cells.

The accumulation of mitochondrial damage, and concomitant degradation of function, is associated with both aging and neurodegenerative disease. Mitophagy also appears to be inherently limited in mature neurons (reviewed in (68)), which may explain why this tissue is more sensitive to mitochondrial damage in the first place. Increasingly, it has been hypothesized and demonstrated that increasing mitophagy in these cells may promote better cellular health and aging (reviewed in (68)).

Unfortunately, a relative dearth of compounds appropriate for this purpose is currently available, and identification of new compounds requires a relatively complex screening process, like the whole-organism phenotypic approach demonstrated herein. Although the eight compounds we identified and studied have considerable promise (especially sertraline), substantial additional study is needed to further understand their effects.

## Methods

### *C. elegans* strains and maintenance

Worms were synchronized by hypochlorite isolation of eggs from gravid adults, followed by hatching of eggs in S Basal. 6,000 synchronized L1 larvae were transferred onto 10 cm standard nematode growth medium (NGM) plates seeded with *Escherichia coli* strain OP50 as a food source (88). After transfer, worms were grown at 20°C for 50 hours prior to experiments, or for three days for the next eggs isolation. Young adult worms were used for all assays unless otherwise noted. Strains used in this study include: N2 Bristol (wild-type), NVK90 |*pink-1*(*tm1779*); *houIs001* {*byEx655* [*Ppink-1*::PINK-1::GFP + *Pmyo-2*::mCherry]}| (9), SJ4103 {*zcIs14* [*Pmyo-3*::GFP^mt^]} (89), VK1241 {*vkEx1241* [*Pnhx-2*::mCherry::LGG-1 + *Pmyo-2*::GFP]} (90), TJ356 {*zIs356* [*Pdaf-16*::DAF-16a/b::GFP + *rol-6*(*su1006*)]} (57), CL2166 {*dvIs19* [*Pgst-4*::GFP::NLS]} (91), AY101 {*acIs101* [*Pirg-5*::GFP + *rol-6*(*su1006*)]} (92), PE255 {*feIs5* [*Psur-5*::luciferase::GFP + *rol-6*(*su1006*)]}, GMC101 {*dvIs100* [*Punc-54*::A-beta-1-42::*unc-54* 3’-UTR + *Pmtl-2*::GFP]} (64), GF66 {*dgEx66* [*Pvha-6*::Q82::YFP + *rol-6*(*su1006*)]} (65), and SS104 [*glp-4*(*bn2*)] (93).

### Bacterial strains

Bacterial strains used in this study included *E. coli* OP50, RNAi-competent OP50 (*xu363*), and RNAi-competent *E. coli* HT115 (obtained from the Ahringer RNAi library). Plasmids were isolated, purified, and sequenced prior to transformation into RNAi-competent OP50 (*xu363*). Transformed bacteria were confirmed to contain the plasmid of interest by sequencing as well.

### RNA interference protocol

RNAi-expressing bacteria were cultured and seeded onto NGM plates supplemented with 25 μg/mL carbenicillin and 1 mM IPTG. For double RNAi, bacterial cultures expressing either *vector(RNAi)* or *pink-1(RNAi)* were mixed with sterility-inducing *cdc-25*.*1(RNAi)* with a 1:1 ratio. For experiments with GMC101 strain, 2,000 synchronized L1 larvae were plated onto 6 cm RNAi plates and grown at 20°C for 48 hours prior to use for experiments. For experiments with GF66 strain, 50 gravid hermaphrodites were transferred onto 10 cm *cdc-25*.*1(RNAi)* plates and grown at 20°C for 3 days or until progenies have reached the L4 stage. Sterile progenies were then transferred to treatment plates for the next experimental step.

### Statistical Analysis

RStudio (version 3.6.3) was used to perform statistical analysis. Chi-square test was used to calculate the significance between qualitative variables. One-way analysis of variance (ANOVA) was performed to calculate the significance of a treatment when there were three or more groups in the experimental setting. To follow, Dunnett’s test (R package DescTools, version 0.99.34) was performed to calculate statistical significance or *p* values between each group of the statistically significant experimental results. Student’s *t*-test analysis was performed to calculate the *p* values when comparing two groups in an experimental setting. All statistical test results were indicated in graphs as follows: NS not significant, \**p* < 0.05, \*\**p* < 0.01, and ***p < 0.001. For each of the experiments described below, at least three biological replicates were performed.

### *C. elegans* library screening and compound exposure assays

Synchronized sterile young adult worms were washed from NGM plates seeded with OP50 into a conical tube and rinsed three times. Worms were then sorted into a 384-well plate (∼25 worms/well) for the initial screen or a 96-well plate (∼100 worms/well, half-area) for all other assays. For the initial high-throughput screen, each well in the 384-well plates contained 50 µM of compounds, and the exposure length was 24 h. For other compound/chemical exposure assays, S Basal supplemented with 50 µM PS compounds, 7 mM sodium selenite (Alfa Aesar), 10 µM CCCP (Sigma), 50 µM rotenone (Sigma), 50 µM juglone (Sigma), 100 µM RPW-24, or DMSO (solvent control) was added into the wells of the 96-well plate to a final volume of 100 µL. Worms were imaged with Cytation5 Cell Imaging Multi-Mode Reader (BioTek Instruments) every day for three days for the initial screen or every two hours for twenty four hours for the other assays.

### Imaging and fluorescence quantification

For visualization of the worm reporter strains NVK90, TJ356, CL2166, AY101, and GF66, Cytation5 automated microscope was used. All imaging experiments were performed with identical settings within three biological replicates of each strain. GFP quantifications were performed by using Gen5 3.10 software and CellProfiler. For visualization of the worm reporter strains SJ4103 and VK1241, worms were immobilized by using 1 mM levamisole and then transferred onto 3% agarose pad. Worms were imaged using fluorescence microscope (Zeiss ApoTome.2 Imager.M2, Carl Zeiss, Germany) with a 63x (SJ4103) or 40x (VK1241) objective magnification.

### Fluorescent-dye staining and quantification

Approximately 400 N2 worms (in four wells, each containing ∼100 worms) were treated with PS compounds or corresponding controls for 15 h in 96-well plate. At 14 h of incubation, fluorescent dye with a final concentration of 10 µM for LysoTracker Red, 10 µM for NAO, 4.375 µM for MitoTracker Red, or 3 µM for DHE, were added. For background subtraction, S Basal without any dye was added. Worms were washed three times to remove any remaining compounds or dye before fluorescence measurement was taken via flow vermimetry (COPAS Biosort, Union Biometrica).

### ATP production measurement

A worm strain carrying firefly luciferase gene followed by GFP (PE255) was used for ATP production measurement. Worms were treated with compounds as described above. ATP measurement was carried out according to the published protocol (94). Essentially, at 18 h of incubation, worms were washed three times to remove any remaining compounds. Luminescence buffer was then added, incubated for 3 minutes, and fluorescence (485/20 excitation and 528/20 emission) and luminescence were measured with Cytation5 (BioTek Instruments).

### Oxygen consumption rate measurement

3,000 N2 worms were sorted into each well of a 6-well plate. PS compounds, vehicle control DMSO, or positive control rotenone, and *E. coli* OP50 (final OD_600_: 0.05) were then added into each well to a final concentration of 50 µM. Two wells for each compound were used, totaling to 6,000 worms per condition. Upon 8 h of incubation, worms were collected and transferred into a 15 mL conical. Worms were washed three times to remove residual compounds. Oxygen consumption was measured by using a biological oxygen monitor (YSI 5300) and a Clark-type oxygen electrode (YSI 5301) (Yellow Springs Instrument) at 20°C as previously described (49). Oxygen consumption was recorded continuously for ten minutes.

### Beta-amyloid-induced paralysis and scoring

Synchronous L1 population of *C. elegans* strain GMC101 (95) was reared on *E. coli* OP50 at 20°C for 48 hours to reach the L4 stage. 60 worms were then transferred onto each 35 mm NGM plates containing 250 µM 5-fluoro-2’-deoxyuridine (FUDR) and PS compounds (see **Fig. S2** for final concentrations) or corresponding DMSO control. Worms were then incubated at 25°C to induce paralysis and scored every day for five days. Paralysis was indicated by inability to complete a full sigmoidal body movement spontaneously or following stimulation with a pick. Paralyzed worms were counted and removed from the plate.

### Polyglutamine protein aggregation scoring

50 gravid hermaphrodites of *C. elegans* strain GF66 expressing polyglutamine (*Pvha-6*::Q82::YFP) (65) were reared on *E. coli* OP50 expressing *cdc-25*.*1(RNAi)* and grown at 20°C for 3 days or until progenies have reached the L4 stage. L4 worms were then transferred onto treatment plates as for the beta-amyloid-induced paralysis assay, but kept at 20°C. After 24 hours, worms were transferred into a 96-well plate, washed three times, immobilized with 1 mM levamisole, and imaged with Cytation 5 automated microscope with a 4x objective magnification. The number of aggregates were counted manually for each of the worms.

### *C. elegans* compound toxicity assay

25 synchronized SS104 (*glp-4*) young adult worms were sorted into 384-well plate. Compounds (final concentration: 100 µM, 50 µM, 25 µM, and 12.5 µM) were mixed with *E. coli* OP50 as food source (final OD_600_: 0.05), and then added into each well. Plates were incubated at 25°C for 72 hours. Plates were washed three times and worms were stained with SYTOX™ Orange nucleic acid dye to stain dead worms. After 14 h of incubation with SYTOX™, plates were washed and imaged with Cytation5 automated microscope. CellProfiler software was used to quantify worms’ death.

### Mammalian cell culture and compound toxicity assay

Human fetal glial cells (SVGp12) and prostate epithelial cells (RWPE-1) were purchased from ATCC (Manassas, VA, USA). SVGp12 cells were cultured in minimum essential Eagle medium (ThermoFisher) containing 10% FBS (fetal bovine serum; Corning, Manassas, VA, USA) and RWPE-1 cells were cultured in Defined Keratinocyte SFM (ThermoFisher) with growth supplement at 37°C in a humidified 5% CO_2_ atmosphere. The solution of penicillin-streptomycin (Gibco, Gaithersburg, MD, USA) was used at 1% final concentration.

Fluorescent cell labeling with Hoechst 33342 (ThermoFisher) and propidium iodide (ThermoFisher) with subsequent automated cell counting was used as cytotoxicity assay as previously described (96). Cytation5 Cell Imaging Multi-Mode Reader with DAPI and Texas Red filter sets (BioTek Instruments) and Gen5 3.10 software were used for imaging and cell counting pipeline.

For each cytotoxicity experiment, cells were seeded at a density of 10^3^ cells/well in 96-well plates and cultured for 24 h prior to the drug treatment. Cells were then treated with PS compounds or DMSO (solvent control) at specified concentrations (see **Fig. 7**) for 72 h in 100 µL of complete media. All viability rates were normalized to the corresponding solvent-control wells. The DMSO concentrations in the incubation mixtures or solvent-control wells never exceeded 0.5% (v/v).

## Acknowledgements

*C. elegans* strains used were obtained from the CGC. We thank Daniel Kirienko for comments on the manuscript and Maria Hancu for technical assistance.

## Competing interests

The authors have declared that no competing interests exist.

## Figure Legends

**Table 1.** Summary of PS compounds’ effects on various mitochondrial parameters and other cellular pathways. NS not significant.

|  | PS30 | PS34 | PS83 | PS103 | PS106 | PS127 | PS135 | PS143 |
| --- | --- | --- | --- | --- | --- | --- | --- | --- |
| Mt fragmentation | High | High | Low | Mid | Mid | High | Mid | High |
| Autophagosome formation | High | Less | Mid | Mid | Mid | Less | Mid | Less |
| Autophagolysosome formation | NS | High | NS | NS | NS | High | NS | High |
| ATP depletion | High | High | NS | NS | NS | High | High | High |
| O<sub>2</sub> consumption rate reduction | NS | High | NS | NS | NS | High | NS | NS |
| Mt mass reduction | NS | High | NS | NS | NS | High | NS | NS |
| Mt ΔΨ reduction | High | NS | NS | High | Mid | NS | High | NS |
| ROS formation | NS | NS | NS | NS | NS | NS | NS | NS |
| GST-4 (SKN-1/Nrf2) induction | NS | Mid | Mid | NS | NS | Mid | NS | Mid |
| DAF-16/FOXO nuclear loc. | NS | High | High | NS | NS | High | NS | High |
| IRG-5 (immune response) | High | NS | NS | NS | NS | NS | High | High |

## Supporting Information

**Figure S1. Chemical structures of the eight PS compounds**.

**Figure S2. Three PS compounds reduced the rate of paralysis in *C. elegans* Alzheimer’s model**. Percent paralysis of *C. elegans* GMC101 that expresses full-length human beta-amyloid upon treatment with **(a-h)** DMSO control compared to **(a)** PS30, **(b)** PS34, **(c)** PS83, **(d)** PS103, **(e)** PS106, **(f)** PS127, **(g)** PS135, and **(h)** PS143. Concentrations of PS compounds were indicated on the graphs. At least three biological replicates with ∼180 worms/replicate were analyzed. *p* values were determined from Student’s *t*-test. NS not significant, \**p* < 0.05, ** *p* < 0.01, *** *p* < 0.001.

**Table S1. Chemical information of the eight PS compounds**.

**Table S2. Tanimoto coefficient of the eight PS compounds as compared to each other**.

## Notes

### Competing Interest Statement

The authors have declared no competing interest.

