## Supplementary figures and images for "Small Molecule Stabilization of PINK-1/PINK1 Improves Neurodegenerative Disease"

### Supplemental Figure 1

**PS30**

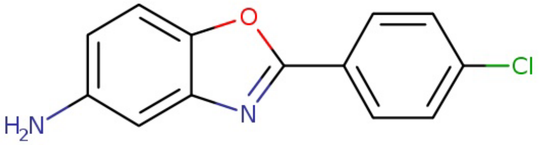

**PS34**

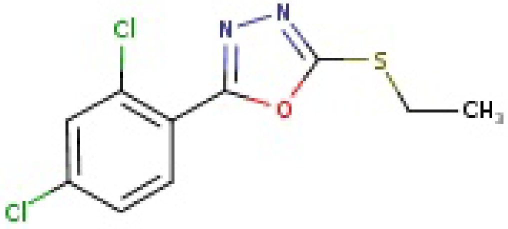

**PS83**

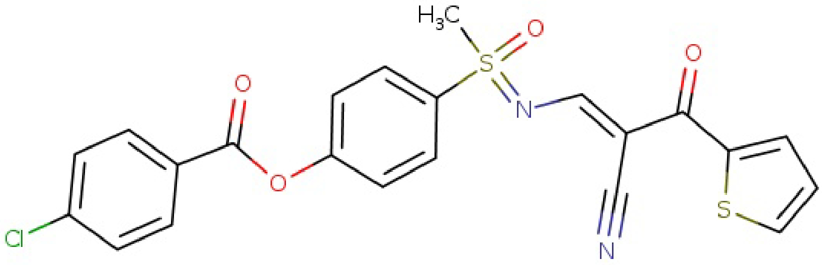

**PS103**

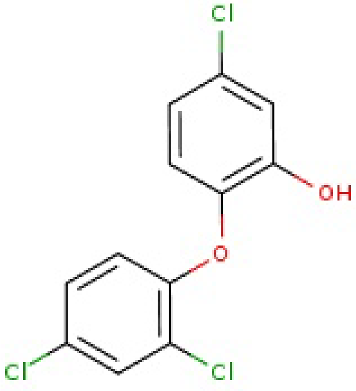

**PS106**

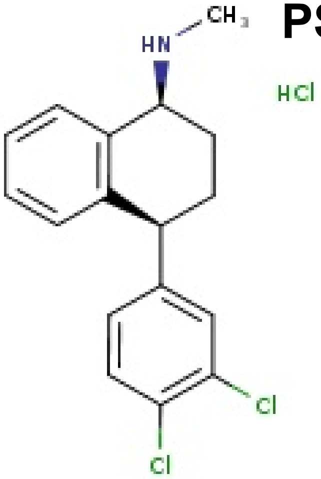

**PS127**

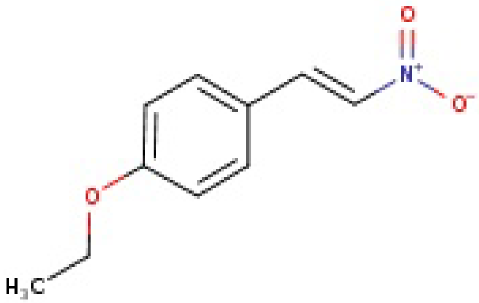

**PS135**

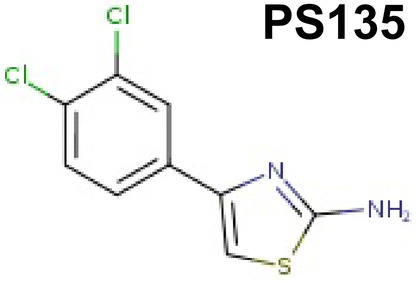

**PS143**

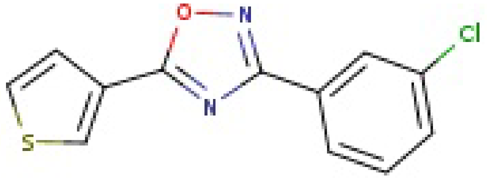

**Figure S1**

### Supplemental Figure 2

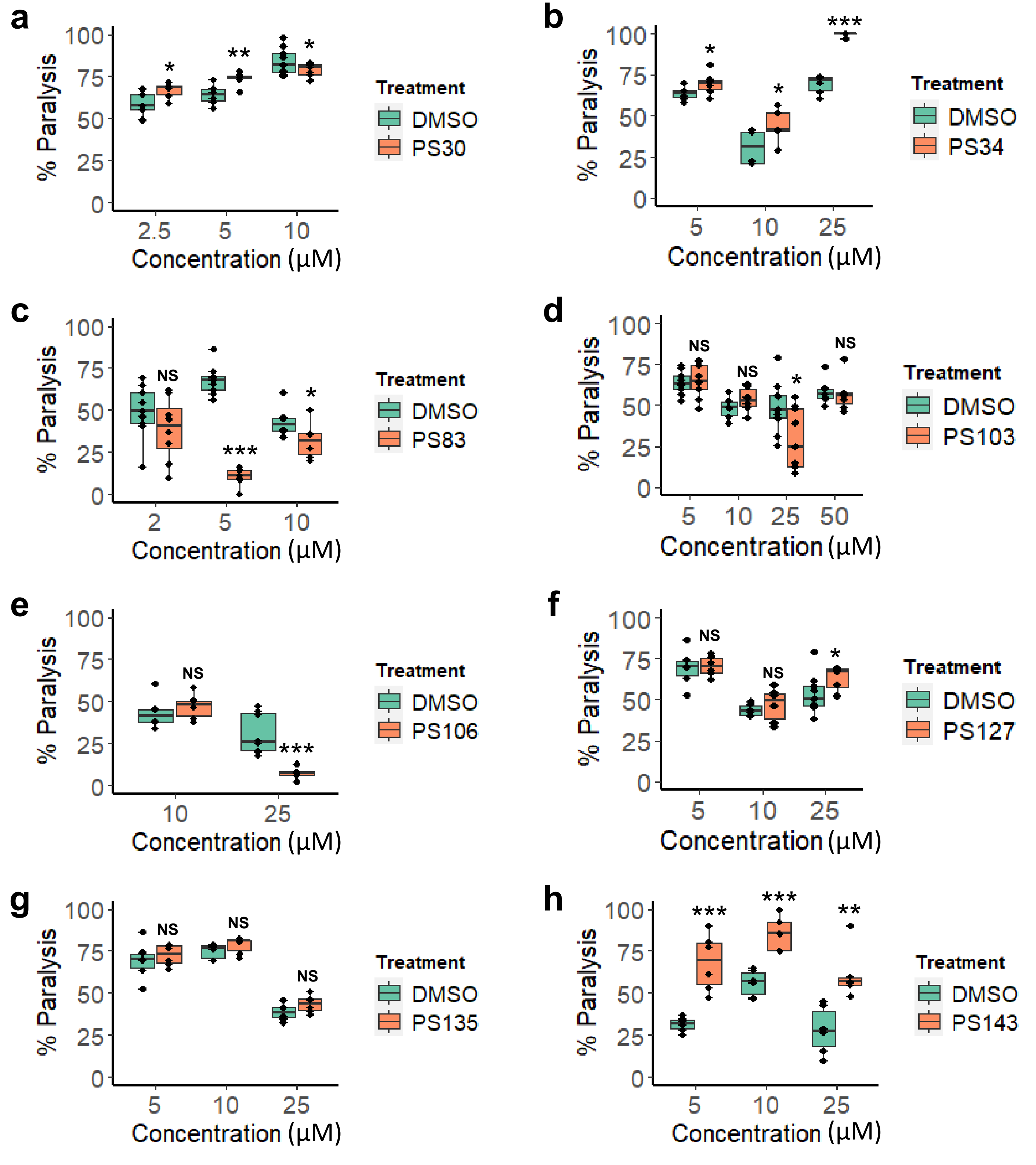
