## Supplemental Table 1 for "Small Molecule Stabilization of PINK-1/PINK1 Improves Neurodegenerative Disease"

| Compound | Library Source | MW (Da) | LogP | Polar area | Donor | Acceptor | Bonds |
|---|---|---|---|---|---|---|---|
| **PS30** | ChemBridge | 244.67 | 3.4 | 52.0 | 1 | 3 | 1 |
| **PS34** | ChemBridge | 275.15 | 3.9 | 64.2 | 0 | 4 | 3 |
| **PS83** | Maybridge | 471.00 | 5.5 | 133.0 | 0 | 7 | 7 |
| **PS103** | NIHCC | 289.50 | 5.0 | 29.5 | 1 | 2 | 2 |
| **PS106** | UT_Kinase | 342.70 | 5.1 | 12.0 | 2 | 1 | 2 |
| **PS127** | ChemBridge | 193.20 | 2.6 | 55.0 | 0 | 3 | 3 |
| **PS135** | ChemBridge | 245.13 | 3.5 | 67.2 | 1 | 3 | 1 |
| **PS143** | Maybridge | 262.72 | 3.8 | 67.2 | 0 | 4 | 2 |

**Table S1**
