## Supplemental Table 2 for "Small Molecule Stabilization of PINK-1/PINK1 Improves Neurodegenerative Disease"

| | PS30 | PS34 | PS83 | PS103 | PS106 | PS127 | PS135 | PS143 |
|---|---|---|---|---|---|---|---|---|
| **PS30** | 1 | | | | | | | |
| **PS34** | 0.291 | 1 | | | | | | |
| **PS83** | 0.134 | 0.158 | 1 | | | | | |
| **PS103** | 0.104 | 0.139 | 0.144 | 1 | | | | |
| **PS106** | 0.192 | 0.167 | 0.157 | 0.135 | 1 | | | |
| **PS127** | 0.119 | 0.126 | 0.183 | 0.218 | 0.110 | 1 | | |
| **PS135** | 0.218 | 0.241 | 0.175 | 0.128 | 0.195 | 0.115 | 1 | |
| **PS143** | 0.192 | 0.184 | 0.191 | 0.096 | 0.178 | 0.102 | 0.267 | 1 |

**Table S2**
